# Acute stress modulates early visual perception and decision-making speed in bees without compromising accuracy

**DOI:** 10.1101/2025.07.31.667877

**Authors:** Olga Procenko, Vivek Nityananda

**Affiliations:** Biosciences Institute, Newcastle University, Henry Wellcome Building, Framlington Place, Newcastle upon Tyne NE2 4HH, UK; School of Biosciences, University of Birmingham, Birmingham B15 2TT, UK

**Keywords:** visual acuity, vision, perception, stress, bees, contrast sensitivity, spatial frequency, decision-making

## Abstract

Acute stress is known to influence decision-making, but it can also directly affect earlier stages of sensory processing. In humans, stress alters the perception of basic visual features, fine-tuning vision to support faster responses in threatening contexts. Whether stress induces similar changes in early vision in insects and thus affects visually guided decision-making, remains unknown. Here, we examined how acute stress, simulated via vigorous shaking resembling a predatory attack, affects bumblebees’ sensitivity thresholds for resolution and contrast sensitivity, two basic visual features, as well as their choice behaviour and decision speed in a dual-choice discrimination task. We found that, similar to humans, stress modulated both thresholds. It increased contrast sensitivity thresholds and shifted spatial frequency thresholds toward finer resolution as well as affecting bee final choice behaviour. Not only were stressed bees more likely to commit to a choice, but they did so significantly faster when their early perceptual decisions were accurate. Collectively, our results suggest that stress reshapes bee visual sensitivity and decision-making not only through an increase in reactivity, but with a flexible shift in decision-making driven by perceptual confidence.

## Introduction

Visually guided behaviour depends on both external stimuli and an animal’s internal state. In insects, a growing body of research shows that vision is modulated by a range of states. For instance, in food deprived blowflies, walking-induced enhancement of motion interneurons is reduced, possibly to conserve energy when reserves are low [1]. Similarly, increased insulin levels in praying mantises—often associated with satiety—reduce the range of distances and azimuths at which prey is detected [2]. Arousal in courting *Drosophila* males has been shown to modulate the gain of specific visual projection neurons, increasing their sensitivity to moving objects and enhancing male persistence while courting a female [3]. Other studies have observed visual modulation associated with internal states such as locomotion [4,5] and arousal [5]. However, the effect of stress on insect vision has remained understudied.

In mammals, the effects of stress-induced states on vision have been extensively studied, including in humans and primates. Stressed participants not only detect threats more quickly [6,7] but also demonstrate enhanced goal-directed visual search performance, even for threat-unrelated objects [8,9]. Being in a high-vigilance state could therefore enhance the visual system’s ability to detect relevant environmental information [10,11].

Fear can also modulate early visual perception, heighten stimulus-driven attention, and, consequently, facilitate visual search. For instance, the brief presentation of a fearful face facilitates target detection by enhancing contrast sensitivity [12]. Fear also leads to increased sensitivity to low-level spatial frequencies, another crucial component of visual acuity [13,14]. Similarly, emotional arousal leads to a shift in peak contrast sensitivity towards lower spatial frequencies [15]. These changes to early vision, therefore, directly influence visual search and suggest an adaptive role in threat detection.

Given the possible adaptive role for these processes, we should expect similar modulation of vision by stress even in insects. Pollinators such as bumblebees, rely heavily on vision for navigation and foraging. They are also exposed to multiple stressors, including natural predators and human-caused environmental challenges. While stress has been shown to alter bee visual decision-making [16], its effects on visual processing remain unexplored. To address this, we investigated whether exposing bees to task-irrelevant stress, mimicking predatory attacks known to induce affective states in bees [16], influences their visual perception. Specifically, we examined if this stress modulates the perception of low-level visual features, stimulus contrast and spatial frequency. Bees were trained to discriminate rewarding horizontal sigmoidal gratings from unrewarded vertical ones in a Y-maze set-up. Subsequently, bees were tested on pairs of gratings with varying spatial frequencies and decreasing contrasts to determine their spatial resolution and contrast thresholds. We hypothesised that exposing bees to simulated predatory attacks, and the resultant negative states would lead to a shift in contrast and spatial resolution thresholds.

## Materials and methods

### Animals and housing

We used four commercially raised *Bombus terrestris* colonies (Agralan, UK) that were transferred to bipartite plastic nest-boxes (28.0 × 16.0 × 12.0 cm). The nest-box was connected to the custom-built Y-maze via a transparent acrylic tunnel (56.0 × 5.0 × 5.0 cm), with shutters controlling bee access to the maze. Bees were kept at 23 ± 2 °C, with double LED tubes (Philips CorePro, 600 mm, 8W T8) providing uniform illumination in the Y-maze: 1100 lux in both arms and 1020 lux at the entrance (measured by digital light meter, Dr. meter LX1010BS, USA). Through the experimental period, colonies were fed with ∼3 g commercial pollen daily (Koppert B. V., The Netherlands) and provided with sucrose solution (20% w/w) *ad libitum* with a gravity feeder on non-experimental days. Bees were food-deprived prior to training or testing. Although invertebrates do not fall under the Animals (Scientific Procedures) Act, 1986 (ASPA), the experimental design and protocols were developed incorporating the 3Rs principles. Housing, maintenance, and experimental procedures were non-invasive and were kept as close as possible to the natural living conditions of the animals. This project was approved by Newcastle University’s ethics board with the reference 8666/2020.

### Training Apparatus

The Y-maze design followed the specifications of a previously published experiment designed for measuring visual acuity in *Bombus terrestris* [17]. The Y-maze comprised three identical arms, each measuring 20 cm in height, width, and length, and covered with UV-transparent Plexiglas sheets (see Fig. 1A). Two of these three arms served as decision arms, while the third functioned as the entrance arm. The entrance arm contained a transparent Plexiglass tunnel (3 cm x 3 cm x 3 cm), which connected to the other two arms within the maze. This tunnel in the entrance arm provided direct access to an outer tunnel leading to the nest-box. The rear wall of each decision arm was equipped with a hole (1cm in diameter) used for the insertion of a reward chamber.

**Figure 1.**
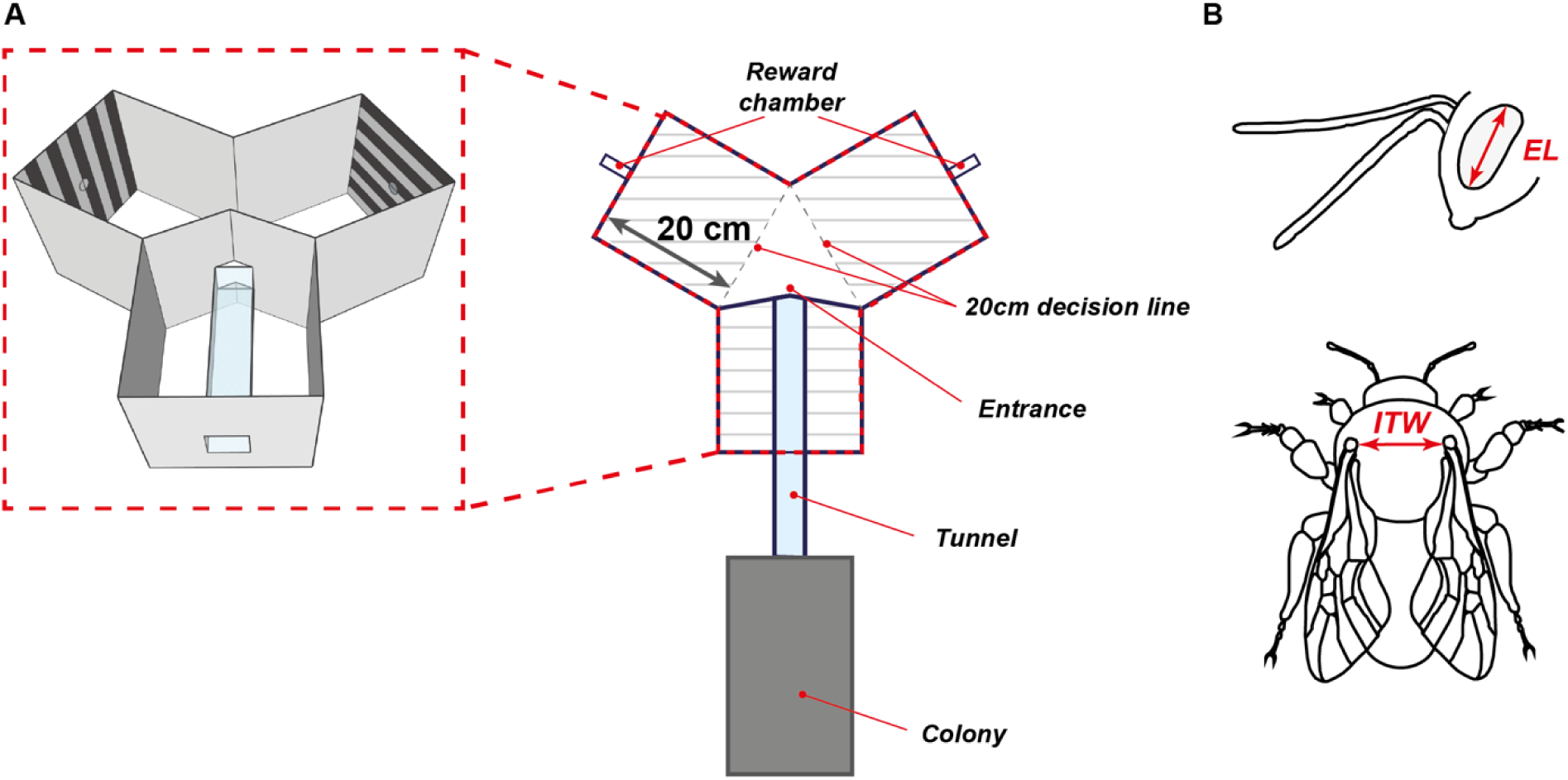
Experimental setup and morphological measurements. **(A)** Experimental setup and stimuli. Bees accessed the decision arms through the entrance at the end of the tunnel. Both decision arms (20 x 20 x 20 cm) contained openings that led to a rewarding chamber which bees had to enter in order to sample its content. During training and testing trials, achromatic sinusoidal gratings were positioned on the back wall of each decision arm. The choice was determined by the first crossing of the 20 cm decision line in either of the two decision arms. **(B)** Morphological measurements included the length of the compound eye (EL) and the intertegular width (ITW).

### Stimuli

Stimuli were identical to those used in a previous study [17], and consisted of achromatic sinusoidal gratings with pattern wavelengths of 6.6, 5, 2.5, 1.53, 1.25, 1.0, and 0.5 cm, corresponding to spatial frequencies of 0.035, 0.070, 0.140, 0.280, 0.437, and 0.699 cycles deg^−1^ of visual angle at the 20 cm decision line (see Fig. 1A). Patterns with spatial frequency of 0.070 cycles deg^−1^ and 87% contrast were used for training because they have previously been shown to be the easiest for bees to resolve [17]. For the contrast sensitivity experiments, we used gratings with a fixed spatial frequency of 0.070 cycles deg^−1^, with varying Michelson contrasts of 89%, 68%, 54%, 39%, and 22%. For the spatial resolution experiment, we used gratings of varying spatial frequencies of 0.035, 0.070, 0.140, 0.280, 0.437, and 0.699 cycles deg^−1^ at a fixed Michelsen contrast of 87%.

### Training and testing procedure

#### Pre-training

Bees foraging on the feeder were marked with a paint marker (Edding 750, Japan) and later recruited for training. To familiarise individual bees with the two reward chambers, each bee was captured at the maze entrance using a cup and guided into a chamber by aligning it with the entrance, forcing the bee to enter and discover a 0.2 ml droplet of 50% (w/w) sugar solution. This procedure was repeated twice for each reward chamber for a total of four trials. During pre-training, bees were presented with conditioned grating pair (spatial frequency of 0.070 cycles deg^−1^ and 87% contrast) for the first time: one positioned horizontally and the other vertically, with one grating in each arm (see Fig. 1A). Through the experiment, the horizontal grating was always associated with a reward, and the vertical grating with water. During pre-training, the initial presentation of the rewarding horizontal grating was randomised across bees. Subsequently, the side of this grating in the three following pre-training trials was alternated based on the initial presentation sequence (e.g., L-R-L-R or R-L-R-L).

#### Training

After the forced-choice pre-training, we began the training phase where bees were allowed to access the reward chamber freely. As before, on each training trial bees were presented with horizontal and vertical gratings positioned in the decision arms. The reward chamber on the arm with the horizontal grating contained a sugar reward (0.2 ml, 50% w/w), while the reward chamber on the side of the vertical grating contained distilled water (0.2 ml). The side of the rewarding horizontal gratings was pseudorandomised across trials, with no more than two consecutive presentations of horizontal gratings on the same side. A new pseudorandom sequence was generated for each bee in R by using the *sample()* function. During training and testing, reward chambers were changed each trial to provide fresh sucrose/distilled water and remove scent marks. Chambers were washed with 70% ethanol and hot water, then dried before the next experimental day.

In previous visual acuity experiments [17], a stationary point at the entrance to the maze was used as the decision point. However, we observed that some bees walked down instead of taking off immediately upon entering the tunnel. Therefore, this entrance stationary point was considered ineffective as a decision point for the study. To avoid inaccurate measurements and following other studies, e.g., Spaethe and Chittka [18], the choice point was defined as the first crossing of the 20 cm decision line in either of the two decision arms. This adjustment allowed bees to navigate a small triangular area after entering but before making their choice. Bees that successfully learned to distinguish between horizontal and vertical gratings and met the learning criterion (80% accuracy in the last 20 trials) proceeded to the testing phase.

#### Testing

Each bee underwent a total of 14 test trials. Each test trial consisted of the presentation of one vertical and one horizontal grating with identical spatial frequencies and contrast levels. To assess tuning to spatial frequencies, each bee was tested on six distinct spatial frequencies in separate test trials, all at a fixed maximum contrast of 87%. The six spatial frequencies used were 0.035, 0.070, 0.140, 0.280, 0.437, and 0.699 cycles deg^−1^. Of these spatial frequencies, three (0.280, 0.437, and 0.699 cycles deg^−1^) were tested twice for each bee, as they are less effectively resolved by bumblebees [17]. To assess contrast sensitivity, bees were presented with gratings featuring a fixed spatial frequency of 0.070 cycles deg^−1^ but with varying contrasts of 87%, 68%, 54%, 39%, and 22%, in separate trials. As before, the bee’s choice was recorded when it crossed the decision line for the first time. Additionally, the final decision of the bee, i.e., entering a reward chamber, was also recorded. Bees failing to enter either reward chamber within a 120-second period were categorised as “omissions” from making a final choice. All test trials were unrewarded, with both reward chambers containing 0.2 ml of distilled water. To keep bees motivated, a set of refresher trials was performed after each test. Each refresher block consisted of two presentations of the grating pair used for training, with a rewarding horizontal grating placed once in each arm in each of the presentations. Bees that failed to enter the reward chamber in the correct arm, indicated by the horizontal grating, after a maximum of three refresher blocks were eliminated from the test trial. In total, 3 bees (1 control, 2 shaken) were excluded.

### Predatory attack simulation

Prior to testing, individual bees were randomly allocated to one of two groups: predatory attack simulation by shaking (n = 20) or an unmanipulated group that served as a control (n = 20). The bees in the shaken group were individually subjected to 60 seconds of shaking at 1200 rpm using a Vortex-T Genie 2 [16]. Before entering the flight arena, we allowed the bee to enter a custom-made tagging cage softened by sponge to prevent physically harming animals while shaking (40 mm diameter, 7.5 cm length). After entering, the bee was gently nudged down with a soft foam plunger until the distance between the plunger and the bottom of the cage was reduced to ∼3 cm. Once the plunger was secured, the cage with the bee was placed inside the vortex cup head. We then ran the Vortex at 1200 rpm for 60 seconds to shake the bee. After the shaking, we released the bee into the tunnel connected to the Y-maze via an opening on the top of the tunnel. The bee was allowed to enter the Y-maze for testing as soon as it was ready to initiate a foraging bout (up to a maximum time of 60 seconds). Bees in the shaking group experienced the shaking procedure before each test trial. After completing all tests, the bee was sacrificed by freezing and stored in 70% ethanol at −20°C. Morphometric measurements, including the length of the compound eye (EL) and the intertegular width (ITW) (see Fig. 1B), were next taken, as earlier studies suggested object resolution is dependent on body size [18]. The measurements were taken under a dissecting microscope using a digital calliper (RS PRO Digital Calliper, 0.01 mm ± 0.03 mm)

### Video analysis

All test trials were recorded using a Huawei Nexus 6P phone with a 1440 × 2560 pixels resolution, capturing video at 120 frames per second (fps). Subsequently, we used the video analysis program BORIS 7.10.2107 [19] to analyse the recorded videos and extract information about bee choices. Specifically, the following time points were coded during analysis: 1) the bee entering the Y-maze, 2) the bee crossing the decision line, and 3) the bee entering the reward chamber. These coded timepoints were used to determine which arm was entered when the decision line was crossed, and which reward chamber was entered. They were also used to calculate latencies for both decisions.

### Statistical analysis

The hypothesis and statistical analyses of the main experiment were preregistered at AsPredicted (#132314). The data were plotted and analysed using RStudio v.3.2.2 (The R Foundation for Statistical Computing, Vienna, Austria, http://www.r-project.org) and custom-written scripts. To determine the thresholds for contrast sensitivity and spatial resolution, we fit a logistic psychometric function (*quickpsy*, “quickpsy” package [20] to the binomial choice data of the bees. In this model, a choice of 1 indicated crossing the decision line in the arm with the rewarding stimulus (hereafter rewarding arm), while a choice of 0 indicated crossing in the arms with the unrewarding stimulus (hereafter non-rewarding arm). The psychometric function is defined as:

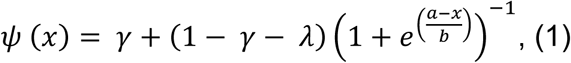

here 𝜓 (𝑥) represents the proportion of correct choices across different spatial frequencies or contrasts. The parameter 𝛾 represents chance performance and was held constant at 0.5, while the parameter 𝜆 accounts for stimulus-independent error or lapse rate, allowing for variability (set as “TRUE”) [20]. The function was fitted using a maximum likelihood method and parametric bootstrapping to estimate confidence intervals.

We fit a generalized mixed effect model (GLMM) with a binomial error distribution and a logit link function (*glmmTMB*, “glmmTBM” package [21] to assess if shaking modulated a bee’s ability to discriminate between horizontal and vertical patterns with decreasing contrasts and varying spatial frequencies. While the effects of spatial frequency and contrast were analysed separately, the response variable in all sets of models was choice accuracy (coded as 1 for crossing the decision line in the correct arm with horizontal grating and 0 for crossing the decision line in the incorrect arm with vertical grating). For the analysis of the effects of contrast, the explanatory variables were Michelson contrast (“Contrast”, coded as a continuous variable and included all five contrasts: 22%, 39%, 54%, 68%, and 87%), treatment (“Treatment”, coded as a factor with two levels: control and shaking) and bee eye width (“Eye width”, coded as a continuous variable). For the analysis of the effect of spatial frequency, the explanatory variables were spatial frequency (“Frequency”, coded as a factor of six levels: 0.699, 0.437, 0.280, 0.140, 0.070, and 0.035 cycles deg^−1^), treatment (“Treatment”: coded as a factor with two levels: control and shaking) and bee eye width (“Eye width”, coded as a continuous variable). In all models, the identity of the bee (“ID”) was included as a random intercept variable.

Both initial choice (crossing the decision line) and final choice (entering the reward chamber) latency were modelled using a linear mixed effect model (LMEM) (*lmer*, lme4 package[22]. As before, the effects of spatial frequency and contrast were analysed separately. Prior to modelling, to handle the right-skewed nature of latency data, we performed log or reciprocal transformations for the initial and final choice latencies respectively. To assess if shaking affected the latency (either initial or final) on the trials with decreasing contrast, the explanatory variables were Michelson contrast (“Contrast”, coded as a factor of five levels: 22%, 39%, 54%, 68%, and 87%), treatment (“Treatment”, coded as a factor with two levels: control and shaking). To analyse the effect of varying spatial frequencies, the explanatory variables were spatial frequency (“Frequency”, coded as a factor of six levels: 0.699, 0.437, 0.280, 0.140, 0.070, and 0.035 cycles deg^−1^), treatment (“Treatment”: coded as a factor with two levels: control and shaking) and bee eye width (“Eye width”, coded as a continuous variable). As before, in all models, the identity of the bee (“ID”) was included as a random intercept variable.

The final choice of reward chamber in each test was categorised according to three mutually exclusive choice categories: choosing a reward chamber on the side of horizontal gratings (correct), choosing a reward chamber on the side of vertical gratings (incorrect), and omission of response (omission). Response omission was recorded if the bee did not enter either reward chamber for longer than the cut-off criterion of 120 seconds. Each choice was transformed into a binary response variable for each choice category. Thus, if a bee entered a reward chamber in the correct arm with horizontal grating, the choice was recorded as (1, 0, 0) for a choice of the correct, incorrect, and omission, respectively. Choices were analysed using generalized mixed linear models (GLMM) using the *glmer* function of the *lme4* package with binomial errors and a logit link function [22]. The independent variables (fixed factors) were the treatment group (Treatment) and the choice category (Choice). As before, the identity of the bee (ID) was included as a random factor.

For each response variable, the model selection process comprised comparing models with and without all possible interactions between explanatory variables. The most appropriate model was selected based on the Akaike information criterion (AIC) scores. We considered the model with the lowest AIC score the best model, i.e., the model that provides a satisfactory explanation of the variation in the data [23]. Models with an AIC difference of less than 2 were considered not significantly better than the model it is being compared to [24]. In such cases, the *anova()* function was used to determine whether adding interaction terms significantly improved model fit. We used the *DHARMa* package [25] for model diagnostics. For each final model, the significance of fixed effects and interactions was assessed using analysis of variance with Wald chi-square tests using the *Anova* function of the *car* package [26]. Additional model summaries were inspected to extract coefficient estimates, standard errors, and associated z- or t-values for fixed effects. If a significant interaction was detected, post hoc comparisons were conducted using the *emmeans* package [27] with Bonferroni adjustment for multiple comparisons

## Results

### Morphometric measurements of shaken and control bees

The intertegular width of the bees ranged from 3.62 mm to 4.27 mm for control bees (median ± IQR: 3.88 ± 0.352 mm) and 3.65 mm to 4.19 mm for shaken bees (median ± IQR: 4.02 ± 0.238 mm). The eye length fluctuated between 2.52 mm to 2.79 mm for control bees (median ± IQR: 2.64 ± 0.142 mm) and 2.42 mm to 2.90 mm for shaken bees (median ± IQR: 2.71 ± 0.137 mm). In line with previous studies (Spaethe and Chittka, 2003), the correlation between these two measures was statistically significant (Control: *r* = 0.75, *p* = 0.0002; Shaking: *r* = 0.70, *p* = 0.0005). While intertegular width did not differ significantly between groups (*W* = 160, *p* = 0.399), eye length showed a moderate difference (*W* = 118, *p* = .044, *r* = .32). Given the strong correlation between the two morphological traits and the significant difference in eye length, the latter was used to control for size-related variation that could explain the observed effects. Consequently, eye length was included as an explanatory factor in subsequent models assessing bee performance.

### Shaken bees have higher contrast thresholds and a shift of spatial resolution threshold towards higher frequencies

#### Contrast thresholds

The contrast threshold was determined by fitting a logistic function to the proportion of correct choices made at the decision line. The data included trials where gratings were expressed at five contrasts (87%, 68%, 54%, 39%, and 22%) with a fixed spatial frequency of 0.070 cycles deg^−1^. A value of 80% correct choices was chosen to measure the threshold for both the control and shaken treatment groups, as this value was exceeded at most contrasts (see Fig. 2A). The control group had a contrast threshold of 22.10% Michelson contrast, while the shaken group had lower contrast sensitivity, with a higher contrast threshold of 35.93%. These results suggest that shaken bees have worse discriminability at lower contrasts, as they require a higher contrast threshold to achieve the same level of performance as the control bees.

**Figure 2.**
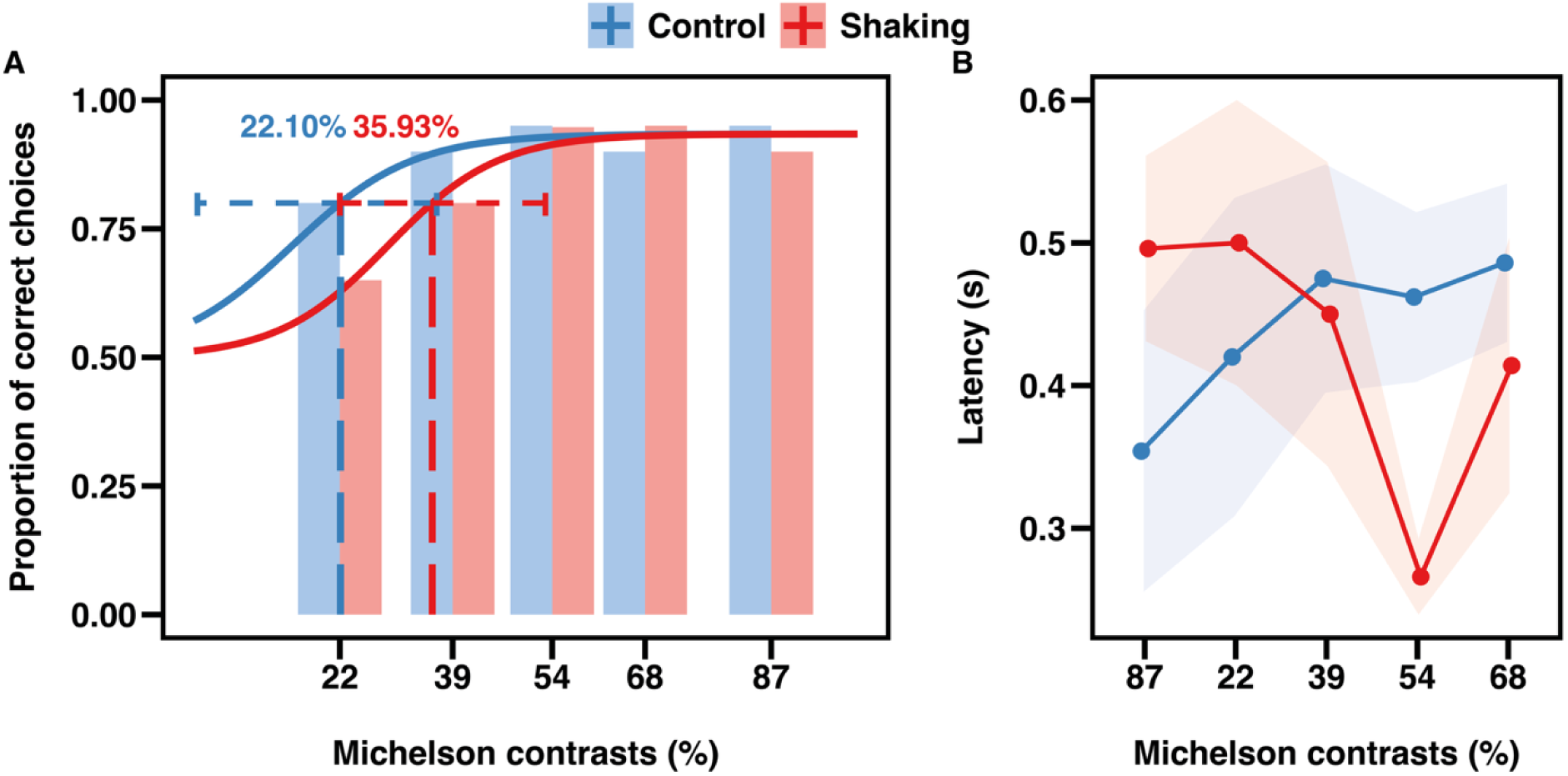
Shaking reduces the contrast sensitivity threshold and choice latency at a contrast of 54%. **(A)** Proportion of correct choices made by control (blue bars) and shaken (red bars) bees at the decision line. The dotted line represents the thresholds at 0.80 proportion of correct choices for both groups, derived from the fitted logistic function (blue solid line: control, red solid line: shaking). Error bars represent the 95% confidence interval for the threshold. **(B)** Average latency to cross the decision line for control (blue) and shaken (red) bees. The averages are presented as a median with the 95% confidence interval (shaded area).

#### Spatial resolution thresholds

To determine the spatial resolution threshold, we fit a logistic function to the proportion of correct choices made at a decision line. Here, data included trials with gratings of varying spatial frequencies (0.035, 0.070, 0.140, 0.280, 0.437, and 0.699 cycles deg^−1^), and fixed contrast (87%). Due to the relatively poor performance, especially at higher spatial frequencies (see Fig. 3A), the value of 65% correct choices were set to measure the threshold for both the control and shaken treatment groups. Although lower than the one used for contrast sensitivity, this value remained significantly above chance (binomial test, *X^2^* = 8.41, *df* = 1, *p* = 0.0037).

**Figure 3.**
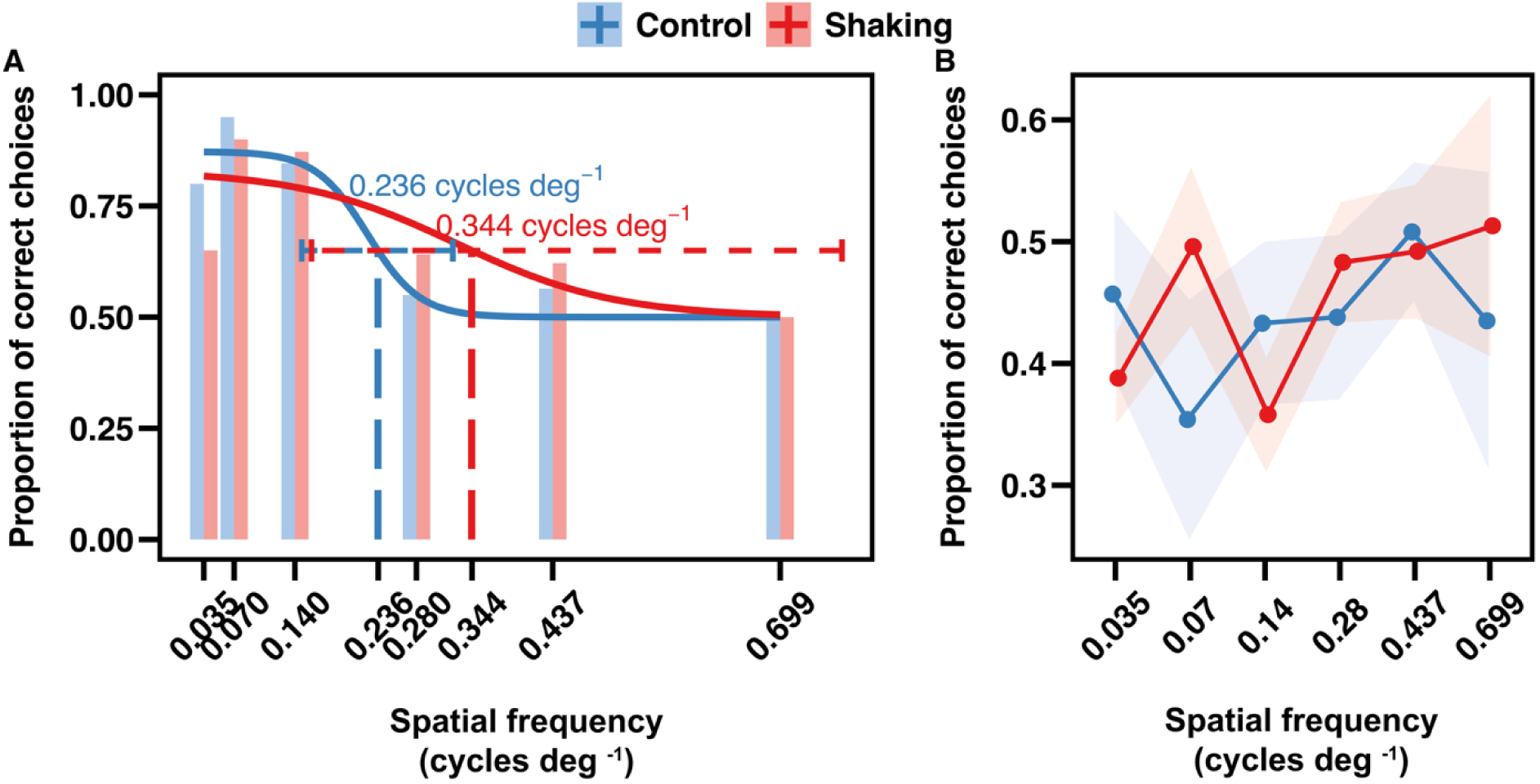
Shaking increases spatial resolution without affecting latency. **(A)** Proportion of correct choices made by control (blue bars) and shaken (red bars) bees at the decision line. The dotted line represents the thresholds at 0.65 proportion of correct choices for both groups, derived from the fitted logistic function (blue solid line control, red solid line shaking). Error bars represent the 95% confidence interval for the threshold. **(B)** Average latency to cross the decision line for control (blue) and shaken (red) bees. The averages are presented as a median with the 95% confidence interval (shaded area).

For control bees, the spatial resolution threshold for discriminating between vertically and horizontally oriented sinusoidal grating patterns was determined to correspond to a spatial frequency of 0.235 cycles deg^−1^. However, the spatial resolution for shaken bees was higher, with a spatial frequency of 0.344 cycles deg^−1^. Therefore, while control bees were able to distinguish gratings expressed in three out of six tested spatial frequencies (0.035, 0.07, 0.140 cycles deg^−1^) with an accuracy above 65%, shaken bees demonstrated the capacity to discriminate patterns expressed at even higher spatial frequencies; including all three that control bees could (0.035, 0.07, 0.140 cycles deg^−1^), in addition to 0.280 cycles deg^−1^. A spatial resolution of 2.91° in shaken bees suggests they can distinguish finer details and patterns at a smaller angular size than the control group.

### Shaking does not affect initial choice accuracy or choice latency at the decision line

#### The effect of contrast sensitivity

Surprisingly, bees demonstrated remarkable accuracy in distinguishing between horizontal and vertical gratings, even when tested at lower contrast levels (see Fig. 2A). Despite the high performance, the contrast threshold for shaken bees was found to be higher (Fig. 2A). We therefore wanted to determine whether this higher contrast threshold had a substantial impact on their overall accuracy in crossing the 20 cm decision line.

The best-fitting model included treatment, contrast, and eye length as fixed predictors without any interaction terms (see Table S1). As contrast levels increased, there was a corresponding increase in the likelihood of bees making the correct choice (Model Estimate ± standard error = 0.030 ± 0.011, *z* = 2.761, *p* < 0.01). Importantly, this trend was consistent across both treatment groups, as shaken bees showed no significant difference in their likelihood of making the correct choice compared to control bees (Model Estimate ± standard error = −0.410 ± 0.505, *z* = −0.811, *p* = 0.417). Eye length also was not a significant predictor of the response accuracy (Model Estimate ± standard error = −0.498 ± 2.309, *z* = −0.022, *p* = 0.8291). This finding demonstrates the ability of bees to maintain discrimination accuracy even as contrast levels decreased, irrespective of their treatment group.

We also explored whether there were variations in the time it took for bees to cross the decision line. The best-fitting model for the choice latency analysis included treatment and contrast as fixed factors and an interaction term (see Table S2). Fixed effects of contrast (*X^2^* = 8.84, *df*: 4, *p* = 0.065) and treatment (*X^2^* = 0.025, *df*: 1, *p* = 0.874) were not significant. However, the effect of treatment was contrast dependent (*X^2^* = 12.69, *df*: 4, *p* = 0.013), with shaken bees making significantly faster choices only at contrast of 54% (*t* = 2.697, *p* = 0.008) (Fig. 2B).

### The effect of spatial frequency

When examining the accuracy of bee performance in crossing the decision line on the trials with varying spatial frequencies, the best-fitting model included treatment, spatial frequency and eye width as fixed predictors without any interaction between factors (see Table S3). Unlike the linear relationship observed with contrast levels (as contrast decreased, accuracy also decreased), the response to varying frequencies displayed a different pattern. The highest accuracy was expected at 0.07 cycles deg^−1^ (conditioned resolution), with a decline in accuracy for both higher and lower frequencies. This was precisely the pattern we observed (Fig. 3A). When keeping a spatial frequency of 0.07 cycles deg^−1^ as a reference group, the performance of bees decreases at the lowest (Frequency 0.035 cycles deg^−1^, Model Estimate ± standard error = −1.480 ± 0.717, *z* = −2.064, *p* = 0.039) as well as the three highest spatial frequencies (Frequency 0.28 cycles deg^−1^, Model Estimate ± standard error = −2.200 ± 0.658, *z* = −3.342, *p* < 0.001; Frequency 0.437 cycles deg^−1^, Model Estimate ± standard error = −2.210 ± 0.660, *z* = −3.347, *p* < 0.001; Frequency 0.699 cycles deg^−1^, Model Estimate ± standard error = −2.682 ± 0.700, *z* = −3.383, *p* < 0.001). At the same time, bees showed no decrease in accuracy when presented with gratings at a spatial frequency of 0.14 cycles deg^−1^, which is double their trained frequency of 0.07 cycles deg^−1^ (Model Estimate ± standard error = −0.723 ± 0.693, *z* = −1.043, *p* = 0.297). This suggests that this lower frequency is well within the bees’ resolvable range and does not impair discrimination. Once again, shaking did not affect the likelihood of crossing the correct decision line (Model Estimate ± standard error = 0.298 ± 0.328, *z* = 0.906, *p* = 0.3650). Similarly, eye width did not significantly affect bee accuracy either (Model Estimate ± standard error = −2.890 ± 1.579, *z* = −1.830, *p* = 0.0672).

We further examined whether the decreased accuracy observed at lower and higher spatial frequencies also led to an increase in latency of crossing the decision line and whether shaking impacted this latency. The best-fitting model for assessing the latency to cross the decision line included treatment and spatial frequency as fixed predictors without considering their interaction (see Table S4). Shaking had no statistically significant effect on crossing latency during trials with varying spatial frequencies (Model Estimate ± standard error = 0.037 ± 0.119, *t* = 0.309, *p* = 0.759). Interestingly, bees took longer to make their choice and cross the decision line at higher spatial frequencies when compared to the trained spatial frequency of 0.07 cycles deg^−1^ (Fig. 3B), which is easily resolvable. Choice latency was significantly higher when the grating had a spatial frequency of 0.699 cycles deg^−1^ (Model Estimate ± standard error = 0.229 ± 0.107, *t* = 2.138, *p* = 0.033). Notably, these were also the spatial frequencies at which bees were less likely to cross decision line in the correct arm (Fig. 3A). Therefore, the increase in choice latency on these trials may indicate a lack of certainty at these highest spatial frequencies.

### The effect of shaking on final choices

Using a two-arm Y-maze allows assessment not only of animals’ early perceptual choices (i.e., initial crossing of the decision line), but also of their subsequent decision-making, as indicated by entry into the reward chamber. Since the arms were not physically separated, bees could freely adjust their flight path and switch to the alternative one if needed. Given the opportunity to gather more information by getting closer to the grating, early perceptual decisions could either be confirmed or corrected. Longer choice latencies have also been linked to lower confidence levels [28]. We therefore measured the time taken to enter the reward chamber, i.e., the final choice latency. To account for cases where bees did not choose even after inspection of both gratings at a close, we set a time limit of 120 seconds. Bees that took longer than this cut-off were considered to be omitting a choice. This approach resulted in three possible decision outcomes - correct choice, incorrect choice, or choice omission. First, we assessed whether stress affects bees’ willingness to make a choice, and thus their likelihood of falling into one of three outcome categories. We then examined whether perceptual accuracy at the initial decision line modulated decision-making speed for the final choice on trials where a choice was made.

#### Fixed spatial frequency and varying contrast

The final model for the likelihood of choices included choice category and treatment as factors (see Table S5). Overall, bees were less likely to make incorrect choices (Model Estimate ± standard error = −3.255 ± 0.269, *z* = −12.084, *p* < 0.0001) or omit from choosing (Model Estimate ± standard error = −6.149 ± 0.736, *z* = −8.363, *p* < 0.0001) compared to making correct choices. The likelihood of making any choice (correct, incorrect, or omission) was also unaffected by shaking (Model Estimate ± standard error = −0.021 ± 0.265, *z* = −0.078, *p* = 0.938) (Fig. 4B).

**Figure 4.**
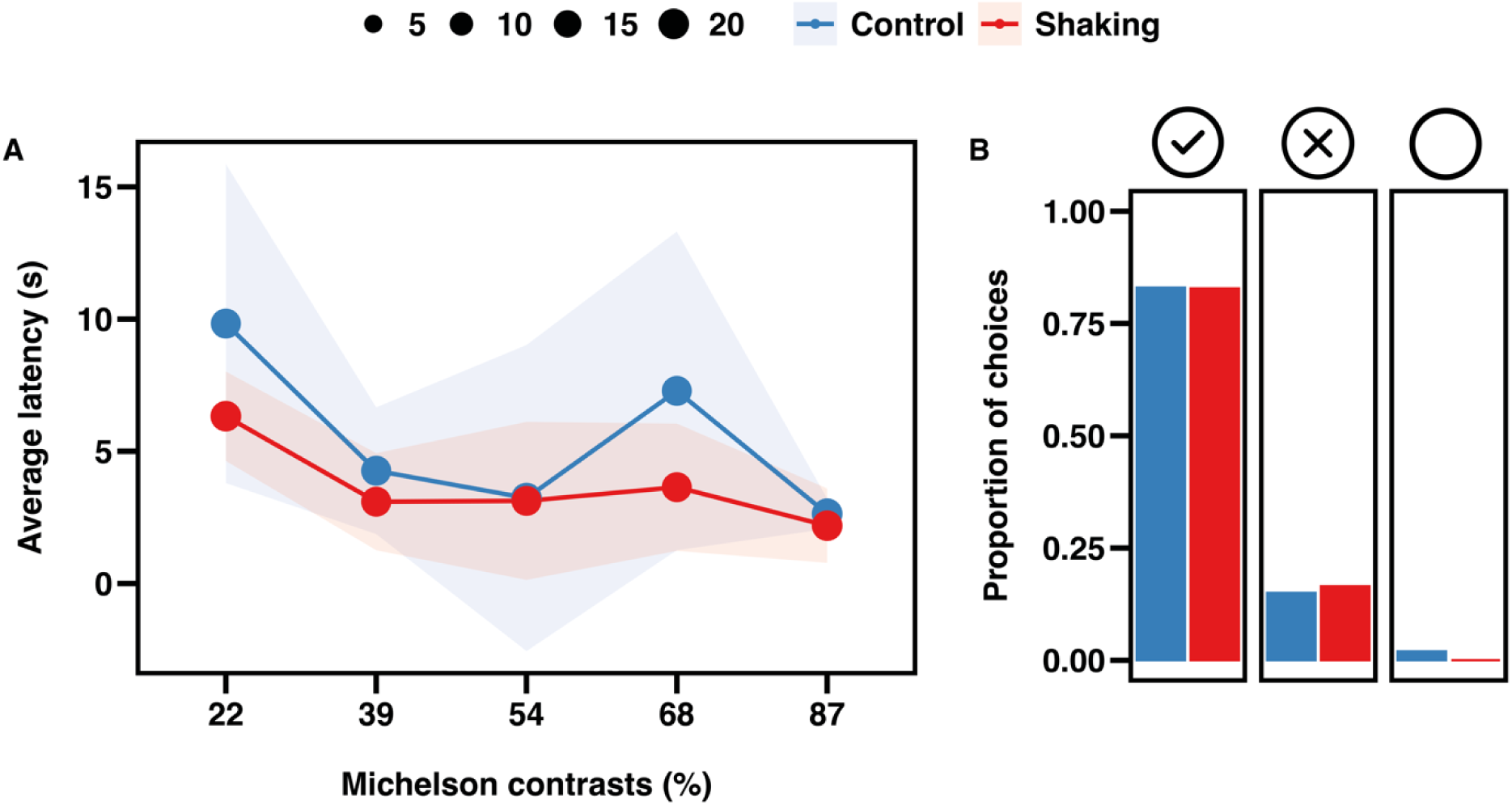
Shaking does not affect the latency of final choices or the probability of choices across contrasts. **(A)** Latency to make correct choices as a function of contrast for Control (blue) and Shaken (red) bees. Dots represent median values with 95% confidence intervals (shaded area). The size of each dot represents the number of bees. **(B)** Proportions of correct choices (left, check mark), incorrect choices (middle, cross mark), and omissions (right, open circle) for Control (blue bars) and Shaken (red bars) bees.

The best-fitting model for assessing final choice latency (entering a reward chamber) included contrast, treatment and accuracy as fixed factors only (see Table S6). As anticipated, with decreasing contrast levels, bees took longer to enter one of the two reward chambers (Model Estimate ± standard error = 0.003 ± 0.001, *t* = 4.830, *p* < 0.001) (Fig. 4A). Interestingly, early perceptual choice accuracy (i.e., crossing the initial decision line in the correct arm) had no significant effect on how long bees took to make their final choice (Model Estimate ± standard error = 0.036 ± 0.041, *t* = 0.886, *p* = 0.377). Additionally, shaking, had a marginally non-significant effect on the final choice latency, despite trending towards a faster decision (Model Estimate ± standard error = 0.069 ± 0.038, *t* = 1.807, *p* = 0.079). Taken together, these results suggest that as stimulus ambiguity decreases with increasing contrast, decision confidence increases. This increase, however, does not further facilitate decision-making. Interestingly, although not statistically significant, stress may contribute to an overall increase in decision speed, yet it had no effect on the likelihood of specific decision outcomes (correct, incorrect, or omission).

#### Fixed contrast and varying spatial frequency

When assessing the likelihood of decision outcome (making correct, incorrect choices, or omissions) the best-fitting model included choice category, treatment, and an interaction between these factors (see Table S7). Unlike with the varying contrast trials, the likelihood of a particular choice depended on whether the bee had been shaken or not (*X^2^* = 9.775, *df*: 2, *p* = 0.008). Shaken bees were overall more likely to enter the correct reward chamber compared to control bees (Model Estimate ± standard error = −0.518 ± 0.261, *z* = –1.986, *p* = 0.047), and less likely to omit responding (Model Estimate ± standard error = 1.218 ± 0.524, *z* = 2.323, *p* = 0.020) (Fig. 6A).

The best-fitting model for the latency data included spatial frequency, treatment and accuracy at decision line as fixed factors, with a significant interaction between treatment and accuracy (see Table S8). When keeping the spatial frequency of 0.07 cycles deg^−1^ (learner resolution) as a reference group, the latency to enter the reward chamber significantly increased for all other spatial frequencies (Frequency 0.035 cycles deg^−1^, Model Estimate ± standard error = 0.782 ± 0.197, *t* = 3.964, *p* < 0.0001; Frequency 0.140 cycles deg^−1^, Model Estimate ± standard error = 1.154 ± 0.170, *t* = 6.798, *p* < 0.0001; Frequency 0.280 cycles deg^−1^, Model Estimate ± standard error = 1.770 ± 0.174, *t* = 10.165, *p* < 0.0001; Frequency 0.437 cycles deg^−1^, Model Estimate ± standard error = 1.915 ± 0.179, *t* = 10.719, *p* < 0.0001; Frequency 0.699 cycles deg^−1^, Model Estimate ± standard error = 2.163 ± 0.212, *t* = 10.218, *p* < 0.0001). The effect of treatment, while not significant alone, was dependent early perceptual decisions (*X^2^* = 7.14, *df*: 1, *p* = 0.008), with shaken bees making significantly faster final choices after crossing the correct decision line (*t* = 4.176, *p* = 0.0001) but not the incorrect one (*t* = 0.044, *p* = 0.965) (Fig. 5).

**Figure 5.**
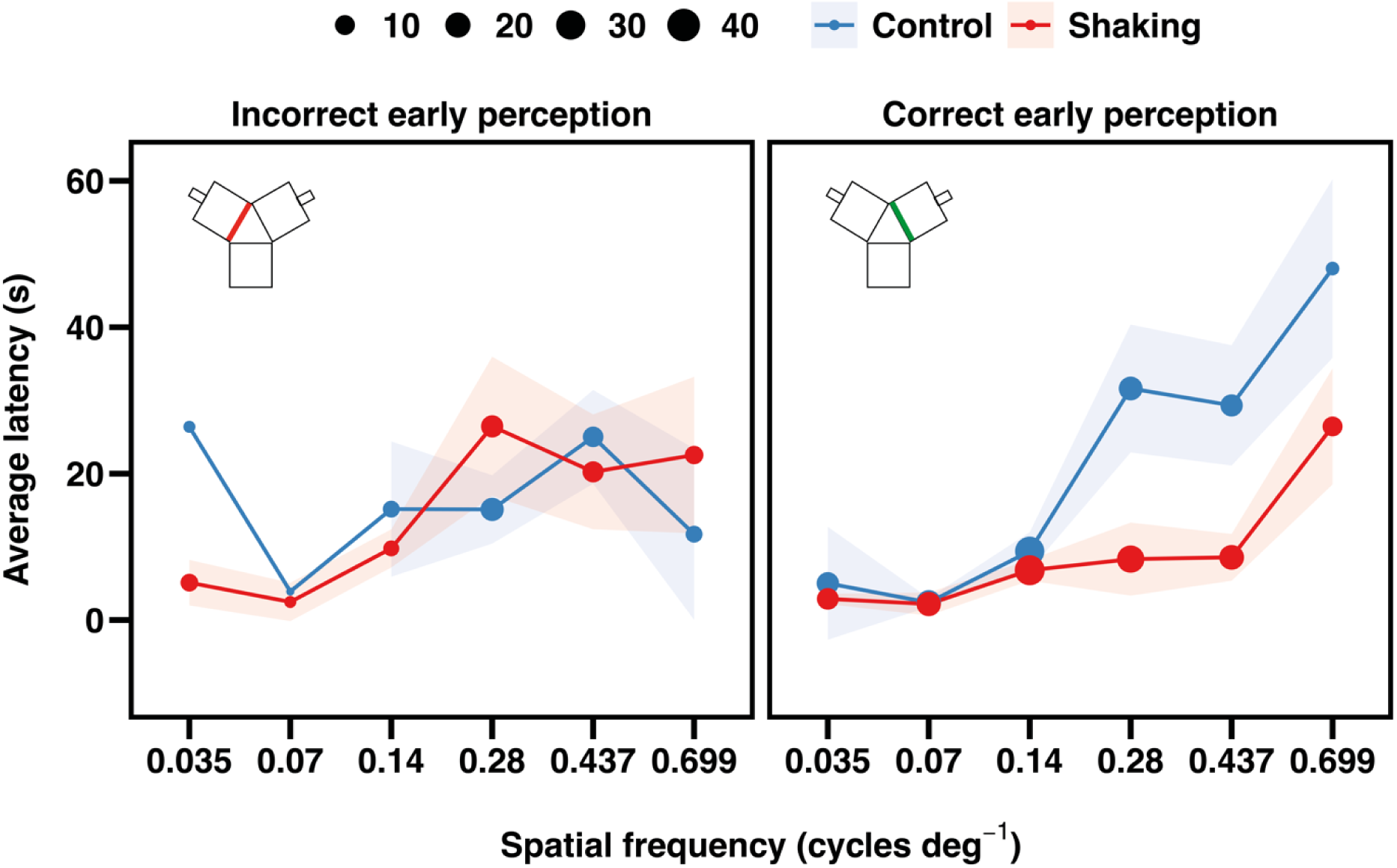
Stress influences speed and outcome of final decisions in the trials with varying spatial frequency. Latency to enter the reward chamber after crossing correct (right panel) or incorrect (left panel) decision point across varying spatial frequencies for Control (blue) and Shaken (red) bees. Dots represent median values with 95% confidence intervals (shaded area). Each dot size represents the number of bees contributing to the median. Note that the graphic depicts a correct choice on the right (green) and an incorrect one on the left (red) but the side of the correct stimulus was counterbalanced across bees.

Faster decisions are not always accurate [29]. We, therefore, next asked whether final choice latencies could predict choice accuracy, and if so, whether or not this was facilitated by stress. The best-fitting model for final choice accuracy data included spatial frequency, choice latency, and treatment as fixed factors, as well as an interaction between choice latency and spatial frequency (see Table S9). While choice latency was a significant predictor (*X^2^* = 5.77, *df*:1, *p* = 0.016) of final choice accuracy, the significant interaction with spatial frequency indicated that its effect varied across spatial frequency conditions (Fig. 6B and Fig. 6C, *X^2^*= 11.14, *df*: 5, *p* = 0.049). Specifically, longer choice durations were significantly associated with decreased accuracy at mid-range spatial frequencies (0.14 cycle deg^-1^: Slope Estimate ± standard error = –0.081 ± 0.029, *p* = 0.0057; 0.28 cycle deg^-1^: Slope Estimate ± standard error = –0.037± 0.017, *p* = 0.032). While not statistically significant, a negative slope was estimated for several other frequencies as well. However, at the finest spatial frequency (0.699 cycle deg^-1^), the trend reversed, showing a non-significant positive association, suggesting that in this condition, longer choice latencies may be associated with increased accuracy (Slope Estimate ± standard error = 0.037± 0.025, *p* = 0.137) (Fig. 6B, 6C). Interestingly, treatment had no additional effect on final choice accuracy (*X^2^* = 0.009, *df*: 1, *p* = 0.924).

**Figure 6.**
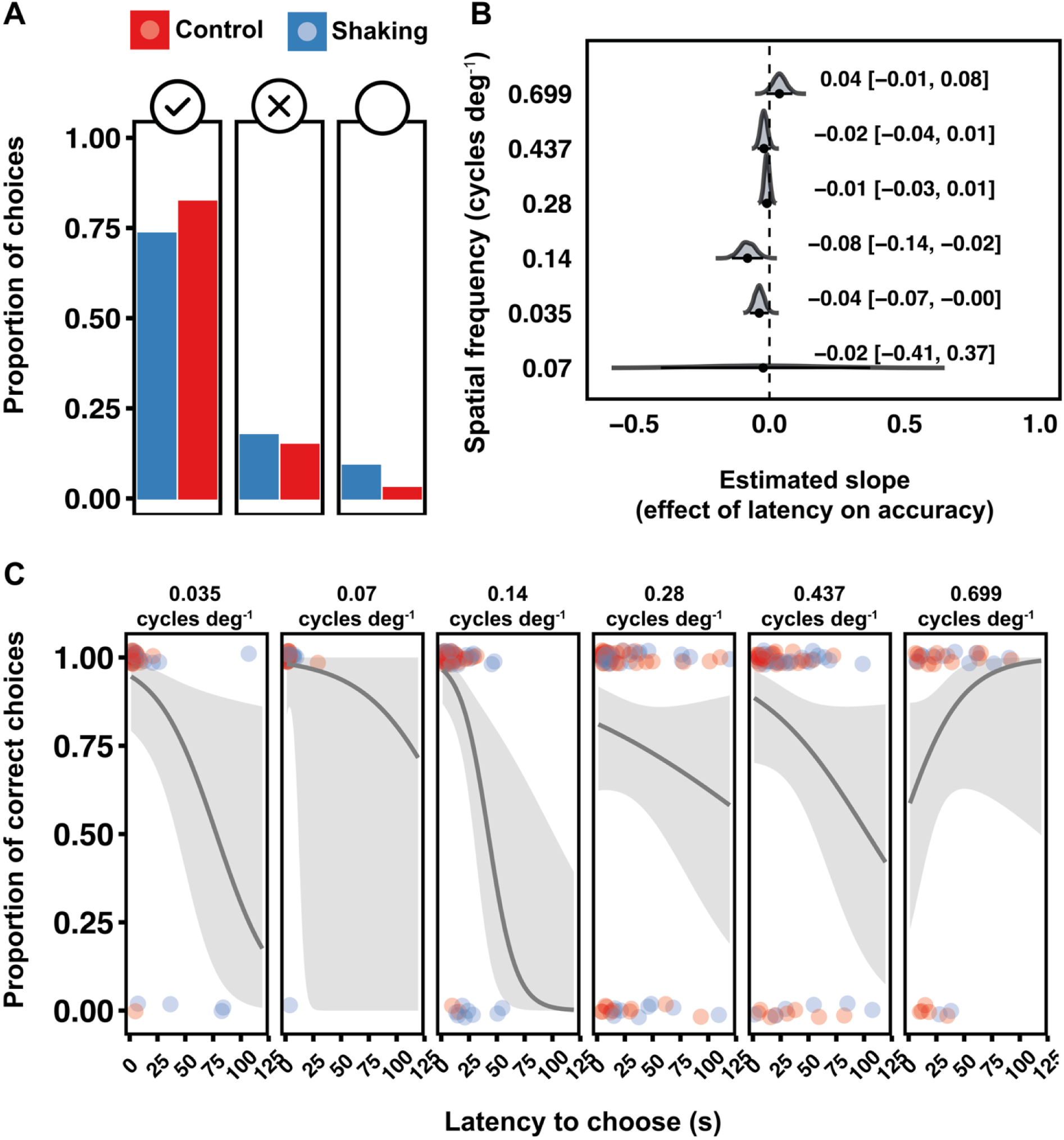
Latency and stress modulate final choice accuracy across spatial frequencies. **(A)** Proportions of correct final choices (left, check mark), incorrect choices (middle, cross mark), and omissions (right, open circle) for Control (blue) and Shaken (red) bees. **(B)** Bootstrapped distributions of slope estimates showing the effect of choice latency on decision accuracy across spatial frequencies. Each distribution represents the uncertainty around the estimated slope; the vertical dashed line indicates no effect. **(C)** Model-predicted relationship between final choice latency and accuracy at each spatial frequency. Coloured points represent observed trial outcomes (1 = correct, 0 = incorrect) for each Control (blue) and Shaken (red) bee. The purple line depicts the predicted probability of a correct choice from the binomial *GLMM*, with shaded 95% confidence intervals. The training stimulus had a spatial frequency of 0.070 cycles deg^-1^.

Taken together, these results suggest that acute stress influences both decision speed and outcome. Shaken bees were less likely to omit making a choice and, when they did choose, responded more quickly—especially following accurate early perception. Interestingly, these faster responses in stressed bees did not result in increased accuracy. While longer choice latencies were associated with reduced accuracy at intermediate spatial frequencies, this relationship was not further modulated by stress.

## Discussion

Internal states like stress influence how visual information is processed to guide behavioural decisions. Here we investigated whether acute stress alters early stages of visual processing in bees — specifically, their sensitivity to fundamental features of spatial vision such as contrast and spatial frequency. Our results demonstrate that exposure to acute stress, shaking that simulates a predatory attack, modulates visual acuity in bees. Stress leads to a reduction in contrast sensitivity and an improved sensitivity to fine spatial details. While these results suggest that, as in mammals, the processing of early visual features in bees can be modulated by stress, the direction of this modulation appears to differ. We find a shift toward enhanced processing of finer visual features in bees and coarser visual features in humans.

In humans, improvements in low-level vision, and consequently, lower contrast thresholds, occur following exposure to stressful threat-inducing stimuli [12]. In addition, the perception of low spatial frequencies is enhanced, and the perception of high spatial frequencies is impaired, with a shift in sensitivity from high to low spatial frequencies [13]. We, however, do not observe this shift in bees – both stressed and control groups maintained peak sensitivity at the trained spatial frequencies (see Fig. 3A); instead, stress flattened the response curve, extending generalisation to higher spatial frequencies, whereas controls showed a steeper decline (see Fig. 3A). While such a phenomenon is better known after absolute conditioning [30,31], threat-related stimuli can also be generalized to conceptually or perceptually similar stimuli [32,33].

Stress-enhanced processing of coarse visual features in humans has been suggested to be adaptive by leading to faster threat detection [34–36]. It might be possible that a similar function is served by the faster processing of finer features in bees. One of the most common predators of bees is the crab spider *Misumena vatia*, which is known to hunt while camouflaged on flowers [37]. To avoid predation, bees typically adopt a side-to-side scanning strategy which, although time-consuming and costly, improves edge detection by gradually amplifying the cryptic spider shape [38]. If acute stress results in a hypervigilant state, perhaps akin to a fear-like condition, it could enhance perception, allowing bees to detect finer features in their surroundings. Therefore, the modulation of visual acuity observed here may indeed facilitate faster processing of threat-related details, serving as an adaptive mechanism that improves the bees’ ability to recognise features of, for example, cryptic predators.

It is also intriguing to compare our threshold estimates from unmanipulated control bees to the findings of the only existing visual acuity study in *Bombus terrestris*. The threshold reported by Chakravarthi et al. (2016)—63.6%—is surprisingly high compared to the contrast threshold estimated here, which is 22%. The experimental design, both Y-maze and stimuli, was consistent across studies. However, the luminance was substantially lower in the previous study (500 lux) compared to ours (1100 lux). Since visual acuity in bees increases logarithmically with increasing light intensity [39], the contrast sensitivity threshold is also expected to change as brightness increases. The estimated spatial frequency threshold in our study, however, closely aligned with the value reported earlier. It is important to note, however, that different accuracy criteria were used to estimate these thresholds: 65% in our study and 75% in Chakravarthi et al. (2016). Interpolating their results to a 65% accuracy criterion yields an estimated threshold of approximately 0.26 cycles deg⁻¹— very close to that observed in the present study. Moreover, bees in our study were conditioned to choose horizontally oriented stimuli, whereas Chakravarthi et al. (2016) used vertical orientations. While orientation could matter, especially given the known differences in interommatidial angle across visual planes in bees [18], this seems unlikely, as a previous study in *Bombus impatiens* reported similar results in a detection task using both horizontal and vertical target orientations [40]. Taken together, the thresholds reported in the current study are largely aligned with earlier estimates for *Bombus terrestris*. Nonetheless, inconsistencies across experimental designs may help explain the observed differences.

While visual acuity thresholds capture early perception of spatial features, the final decision represents a longer-term decision-making process that is driven by increased sampling time and evidence accumulation [41]. In the varying contrast trials, all bees made final choices very quickly (approximately 2.39 s at the highest and 7.27 s at the lowest contrasts; see Fig. 4A). Given the high accuracy and low sensitivity thresholds (see Fig. 2A, 4B), it becomes evident that the selected contrasts were well within the comfortable range of bees’ vision. This, as discussed earlier, is due to the high luminance of our Y-maze, which enabled discriminability between horizontal and vertical patterns even at the lowest contrast. Although shaking trended toward reducing choice latency, it likely could not further enhance response times that were already near ceiling in our setup.

When it comes to varying spatial resolution, however, the effect of shaking was stronger. The range of spatial frequencies in our experiments were much harder to resolve, especially when substantially different from the trained frequency. Given the uncertainty of perceptual evidence, bees became more hesitant when making final choices. This was further confirmed by an overall increased number of omissions and incorrect choices, which was particularly true for control bees. Shaken bees, however, were more likely to commit to choosing, and when committing, responded faster (see Fig. 5). Interestingly, such increased responsiveness appeared to be perception-driven – significantly faster final choices were made only if the early decisions of shaken bees were correct, with this effect becoming most pronounced at the highest spatial frequencies. In spite of the expected speed-accuracy trade-off [29,42], heightened reactivity under stress did not compromise accuracy, as we found no evidence of stress affecting performance quality. Reduced reaction times following acute stress have been reported not only in bees [16], but also in humans [43,44], where the activation of automated responses and suppression of more deliberative decision-making processes suggested as a driving mechanism [44].

Interestingly, however, in humans, if the stressor is mild, such increased reactivity may occur without compromising accuracy suggesting a more nuanced interaction between stress and cognition [45]. Stress effects on human cognition are typically linked to hypothalamo-pituitary-adrenal axis activation. In bees similar stress response pathways have been proposed [46], which may underlie their faster yet accurate responses [47]. Specifically, shaking may have increased arousal via the release of biogenic amines, such as octopamine and dopamine, both of which are known to modulate arousal and attention [46]. This elevated arousal could, in turn, enhance task engagement and reduce choice hesitance, particularly when early perceptual information (i.e., crossing the decision line) was correct. While this remains to be confirmed, biogenic amine-driven arousal may explain stress-induced increases in the willingness to choose, and the faster, yet still accurate, decisions observed in shaken bees.

The effect of stress on visual signal processing are complex. While short term allocation of resources to overcome acute stress tunes visual systems to better detect relevant information, the trade-off of such enhancement on more deliberate decision-making processes may be detrimental. Our study is the first to investigate these interactions in bees–species that rely on vision for decision-making and are continuously exposed to multiple stressors. We show that stress dynamically reshapes how bees process basic visual features and act on visual information, suggesting a broader strategy used to adapt to uncertain environments.

## Acknowledgements

This study was funded a BBSRC David Phillips Fellowship (BB/S009760/1, awarded to V.N.). O.P. was supported by a PhD studentship from the Faculty of Medical Sciences, Newcastle University.

## Author contributions

O.P.: conceptualization, data curation, formal analysis, visualization, writing (original draft preparation). V.N..: funding acquisition, conceptualization, methodology, supervision, resources, writing (review and editing).

## Data availability

Data underlying the figures presented in the manuscript are available at https://figshare.com/s/4fcb82f2e4dcf0e84494

## Supplementary Information

**Table 1S.** Summary and comparison of generalized linear mixed-effects models assessing bee accuracy in crossing a 20 cm decision line on varying contrast trials.

| Selection of best-fit model |  |  |  |  |  |  |
| --- | --- | --- | --- | --- | --- | --- |
| Model | Dependent variable | Fixed factors | Random factor | df | AIC | ΔAIC |
| model 1 | Accuracy | Treatment+Contrast+Eye length | ID | 5 | 149.34 | 0 |
| model 2 | Accuracy | Treatment+Contrast*Eye length | ID | 6 | 150.255 | 0.915 |
| model 3 | Accuracy | Treatment*Eye length+contrast | ID | 6 | 150.883 | 1.542 |
| model 4 | Accuracy | Treatment*Contrast+Eye length | ID | 6 | 150.99 | 1.65 |
| model 5 | Accuracy | Treatment*Contrast*Eye length | ID | 9 | 154.391 | 5.051 |
| Follow-up likelihood ratio test comparing models |  |  |  |  |  |  |
| Model | Parameters | Deviance | X<sup>2</sup> | df | P |  |
| model 1 | 5 | 139.34 |  |  |  |  |
| model 2 | 6 | 138.25 | 1.086 | 1 | 0.298 |  |
| model 3 | 6 | 138.88 | 0 | 0 | 1 |  |
| model 4 | 6 | 138.99 | 0 | 0 | 1 |  |
| Summary of model coefficients with standard errors and significance tests |  |  |  |  |  |  |
| Term | Estimate | SE | z | P |  |  |
| (Intercept) | 2.083 | 6.165 | 0.338 | 0.735 |  |  |
| Treatment Shaking | -0.41 | 0.505 | -0.811 | 0.417 |  |  |
| Contrast | 0.03 | 0.011 | 2.761 | 0.006 |  |  |
| Eye length | -0.498 | 2.309 | -0.216 | 0.829 |  |  |

The tables present the best-fitting generalized linear mixed-effects models, including fixed and random effects, degrees of freedom (df), and Akaike's Information Criterion (AIC). The model with the lowest AIC was selected as best; however, if competing models had a $\Delta AIC < 2$ , they were considered statistically indistinguishable (Burnham & Anderson, 2004), and the simpler model was preferred. In such cases, a follow-up likelihood ratio test was used to compare model fit. The final selected model is shown in italics. $P < 0.05$ indicates statistical significance.

**Table S2.**
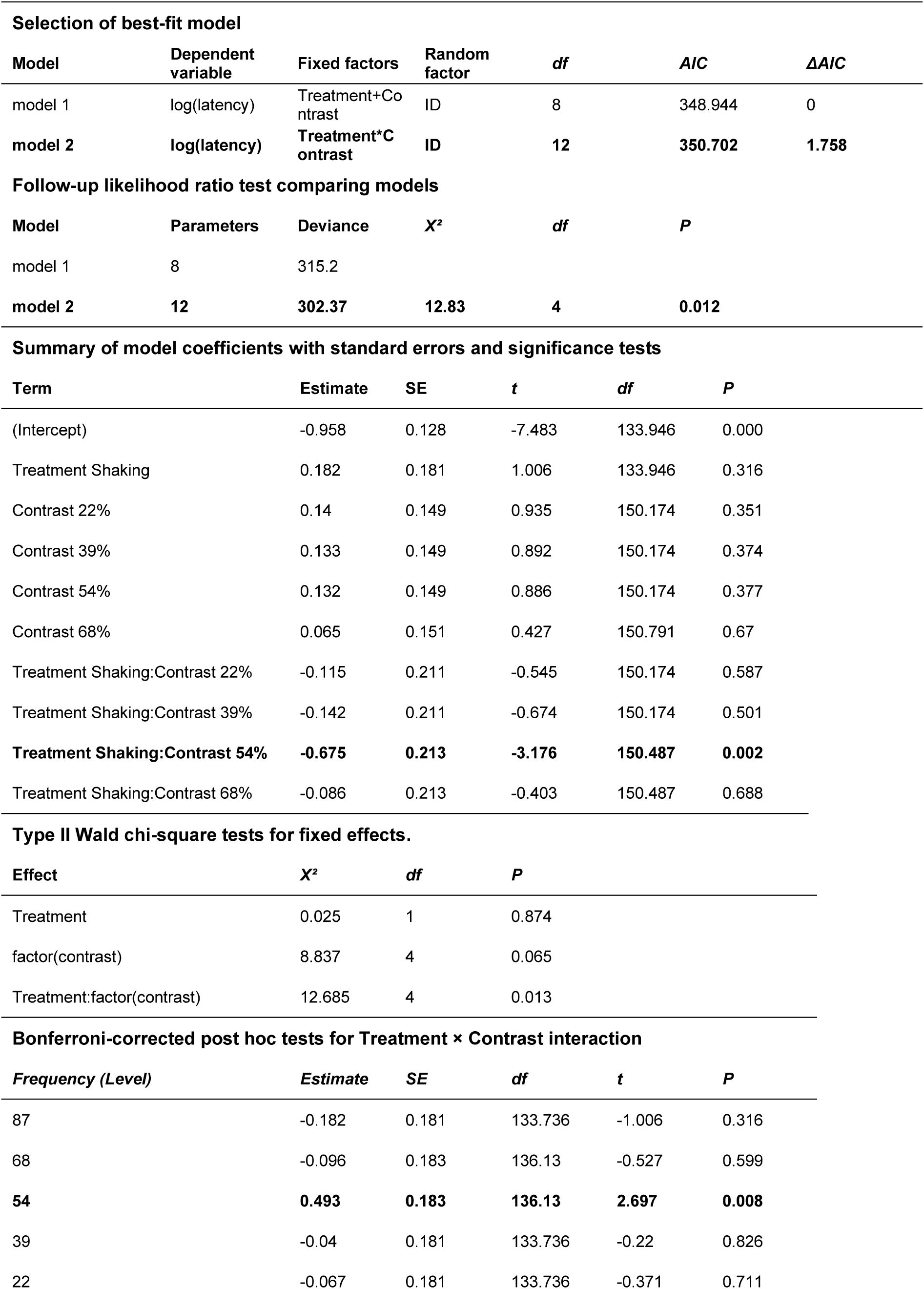
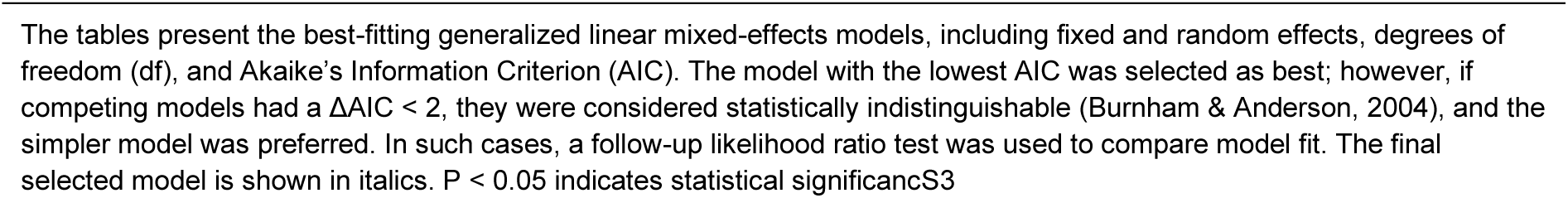
Summary and comparison of generalized linear mixed-effects models assessing bee latency to cross 20 cm decision line on varying contrast trials.

**Table 3S.** Summary and comparison of generalized linear mixed-effects models assessing accuracy in correctly crossing the 20 cm decision line on varying spatial frequency trials.

| Selection of best-fit model |  |  |  |  |  |  |
| --- | --- | --- | --- | --- | --- | --- |
| Model | Dependent variable | Fixed factors | Random factor | df | AIC | ΔAIC |
| model 1 | Accuracy | Treatment+frequency+Eye length | ID | 9 | 396.995 | 0 |
| model 2 | Accuracy | Treatment*Eye length+Frequency | ID | 10 | 397.498 | 0.502 |
| model 3 | Accuracy | Treatment*Frequency+Eye length | ID | 14 | 403.668 | 6.673 |
| model 4 | Accuracy | Treatment+Frequency*Eye length | ID | 14 | 404.174 | 7.178 |
| model 5 | Accuracy | Treatment*Frequency*Eye length | ID | 25 | 418.684 | 21.688 |
| Follow-up likelihood ratio test comparing models |  |  |  |  |  |  |
| Model | Parameters | Deviance | X<sup>2</sup> | df | P |  |
| model 1 | 9 | 379 |  |  |  |  |
| model 2 | 10 | 377.5 | 1.498 | 1 | 0.221 |  |
| Summary of model coefficients with standard errors and significance tests |  |  |  |  |  |  |
| Term | Estimate | SE | z | P |  |  |
| (Intercept) | 10.255 | 4.267 | 2.403 | 0.016 |  |  |
| Treatment Shaking | 0.298 | 0.328 | 0.906 | 0.365 |  |  |
| Frequency 035 deg<sup>-1</sup> | -1.48 | 0.717 | -2.064 | 0.039 |  |  |
| Frequency 0.14 deg<sup>-1</sup> | -0.723 | 0.693 | -1.043 | 0.297 |  |  |
| Frequency 0.28 deg<sup>-1</sup> | -2.2 | 0.658 | -3.342 | 0.001 |  |  |
| Frequency 0.437 deg<sup>-1</sup> | -2.21 | 0.66 | -3.347 | 0.001 |  |  |
| Frequency 0.699 deg<sup>-1</sup> | -2.682 | 0.7 | -3.832 | 0 |  |  |
| Eye length | -2.89 | 1.579 | -1.83 | 0.067 |  |  |
| Type II Wald chi-square tests for fixed effects |  |  |  |  |  |  |
| Effect | X<sup>2</sup> | df | P |  |  |  |
| Treatment | 0.821 | 1 | 0.365 |  |  |  |
| Frequency | 30.543 | 5 | 0.000 |  |  |  |
| Eye length | 3.349 | 1 | 0.067 |  |  |  |

The tables present the best-fitting generalized linear mixed-effects models, including fixed and random effects, degrees of freedom (df), and Akaike's Information Criterion (AIC). The model with the lowest AIC was selected as best; however, if competing models had a $\Delta$ AIC < 2, they were considered statistically indistinguishable (Burnham & Anderson, 2004), and the simpler model was preferred. In such cases, a follow-up likelihood ratio test was used to compare model fit. The final selected model is shown in *italics*. P < 0.05 indicates statistical significance.

**Table 4S.** Summary and comparison of generalized linear mixed-effects models assessing bee latency to cross 20 cm decision line on varying spatial frequency trials.

| Selection of best-fit model |  |  |  |  |  |  |
| --- | --- | --- | --- | --- | --- | --- |
| Model | Dependent variable | Fixed factors | Random factor | df | AIC | $\Delta AIC$ |
| model 1 | log(duration) | Treatment+Frequency | ID | 9 | 584.446 | 0 |
| model 2 | log(duration) | Treatment*Frequency | ID | 14 | 601.475 | 17.029 |
| Summary of model coefficients with standard errors and significance tests |  |  |  |  |  |  |
| Term | Estimate | SE | t | df | P |  |
| (Intercept) | -0.885 | 0.11 | -8.022 | 106.317 | 0 |  |
| Treatment: Shaking | 0.037 | 0.119 | 0.309 | 38.225 | 0.759 |  |
| Frequency 0.035 deg-1 | -0.025 | 0.107 | -0.234 | 308.231 | 0.815 |  |
| Frequency 0.14 deg-1 | -0.048 | 0.093 | -0.514 | 308.354 | 0.608 |  |
| Frequency 0.28 deg-1 | 0.051 | 0.093 | 0.549 | 308.292 | 0.583 |  |
| **Frequency 0.437 deg-1** | **0.183** | **0.094** | **1.952** | **308.467** | **0.052** |  |
| **Frequency 0.699 deg-1** | **0.229** | **0.107** | **2.138** | **308.231** | **0.033** |  |

The tables present the best-fitting generalized linear mixed-effects models, including fixed and random effects, degrees of freedom (df), and Akaike's Information Criterion (AIC). The model with the lowest AIC was selected as best; however, if competing models had a $\Delta AIC < 2$ , they were considered statistically indistinguishable (Burnham & Anderson, 2004), and the simpler model was preferred. In such cases, a follow-up likelihood ratio test was used to compare model fit. The final selected model is shown in italics. $P < 0.05$ indicates statistical significance.

**Table 5S.**
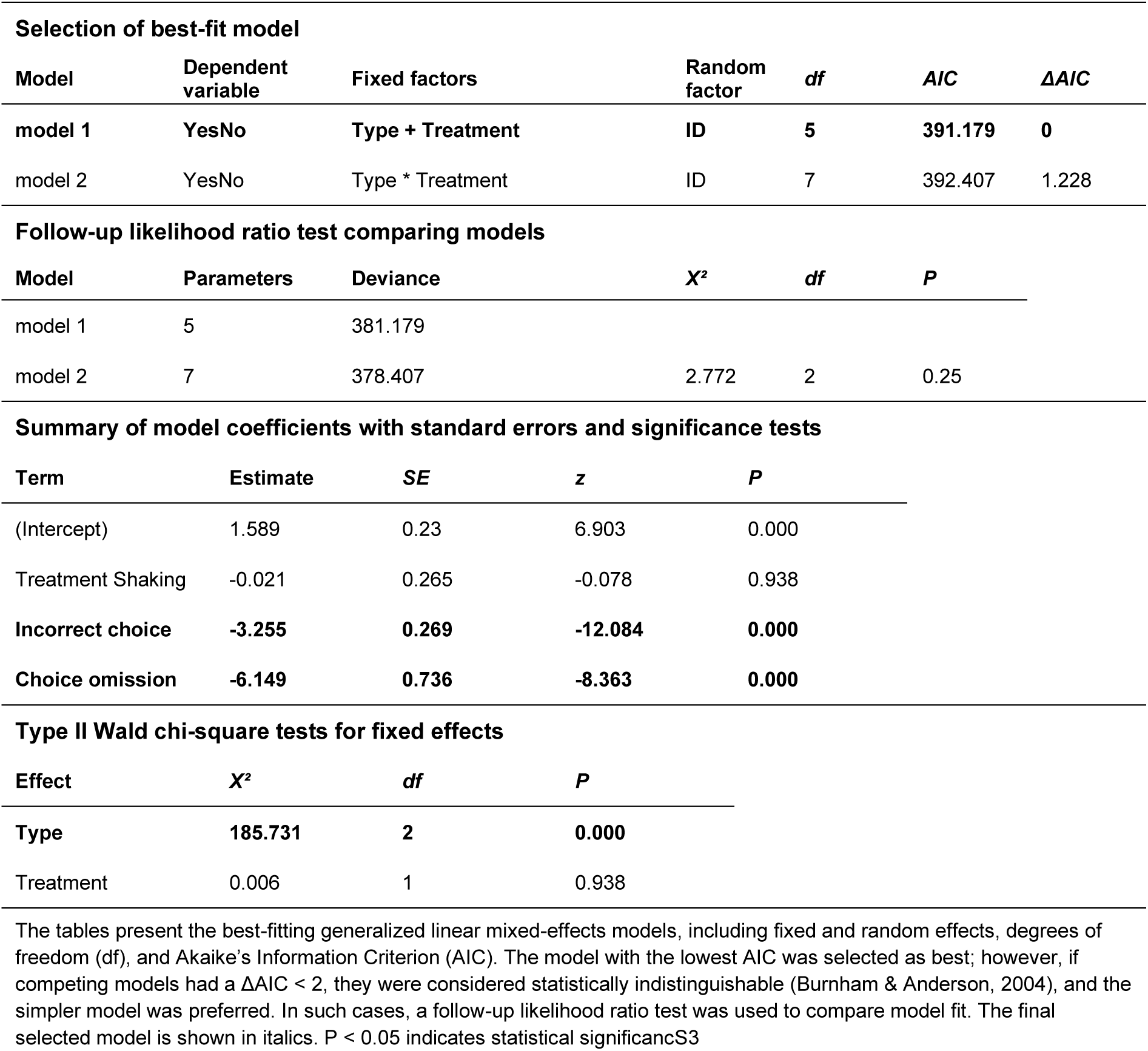
Summary and comparison of generalized linear mixed-effects models assessing bees’ final choice outcomes (correct, incorrect, or omission when choosing which of two rewarding chambers to enter) across varying contrast trials.

| Selection of best-fit model |  |  |  |  |  |  |
| --- | --- | --- | --- | --- | --- | --- |
| Model | Dependent variable | Fixed factors | Random factor | df | AIC | ΔAIC |
| model 1 | YesNo | Type + Treatment | ID | 5 | 391.179 | 0 |
| model 2 | YesNo | Type * Treatment | ID | 7 | 392.407 | 1.228 |
| Follow-up likelihood ratio test comparing models |  |  |  |  |  |  |
| Model | Parameters | Deviance | X² | df | P |  |
| model 1 | 5 | 381.179 |  |  |  |  |
| model 2 | 7 | 378.407 | 2.772 | 2 | 0.25 |  |
| Summary of model coefficients with standard errors and significance tests |  |  |  |  |  |  |
| Term | Estimate | SE | z | P |  |  |
| (Intercept) | 1.589 | 0.23 | 6.903 | 0.000 |  |  |
| Treatment Shaking | -0.021 | 0.265 | -0.078 | 0.938 |  |  |
| Incorrect choice | -3.255 | 0.269 | -12.084 | 0.000 |  |  |
| Choice omission | -6.149 | 0.736 | -8.363 | 0.000 |  |  |
| Type II Wald chi-square tests for fixed effects |  |  |  |  |  |  |
| Effect | X² | df | P |  |  |  |
| Type | 185.731 | 2 | 0.000 |  |  |  |
| Treatment | 0.006 | 1 | 0.938 |  |  |  |

The tables present the best-fitting generalized linear mixed-effects models, including fixed and random effects, degrees of freedom (df), and Akaike's Information Criterion (AIC). The model with the lowest AIC was selected as best; however, if competing models had a $\Delta AIC < 2$ , they were considered statistically indistinguishable (Burnham & Anderson, 2004), and the simpler model was preferred. In such cases, a follow-up likelihood ratio test was used to compare model fit. The final selected model is shown in *italics*. $P < 0.05$ indicates statistical significance.

**Table S6.** Summary and comparison of generalized linear mixed-effects models assessing bees’ final choice latency during varying contrast trials.

| Selection of best-fit model |  |  |  |  |  |  |
| --- | --- | --- | --- | --- | --- | --- |
| Model | Dependent variable | Fixed factors | Random factor | df | AIC | $\Delta AIC$ |
| model 1 | Latency <sup>-1</sup> | Treatment+Contrast+Accuracy | ID | 6 | -68.626 | 0 |
| model 2 | Latency <sup>-1</sup> | Treatment*Accuracy+Contrast | ID | 7 | -63.881 | 4.746 |
| model 3 | Latency <sup>-1</sup> | Treatment+Contrast*Accuracy | ID | 7 | -55.904 | 12.722 |
| model 4 | Latency <sup>-1</sup> | Treatment*Contrast+Accuracy | ID | 7 | -54.909 | 13.717 |
| model 5 | Latency <sup>-1</sup> | Treatment*Contrast*Accuracy | ID | 10 | -26.131 | 42.496 |
| Summary of model coefficients with standard errors and significance tests |  |  |  |  |  |  |
| Term | Estimate | SE | t | df | P |  |
| (Intercept) | 0.077 | 0.05 | 1.548 | 169.141 | 0.124 |  |
| Treatment: Shaking | 0.069 | 0.038 | 1.807 | 38.235 | 0.079 |  |
| Contrast | 0.003 | 0.001 | 4.83 | 154.74 | 0.000 |  |
| Correct choice | 0.036 | 0.041 | 0.886 | 180.986 | 0.377 |  |
| Type II Wald chi-square tests for fixed effects |  |  |  |  |  |  |
| Effect | X<sup>2</sup> | df | P |  |  |  |
| Treatment | 3.264 | 1 | 0.071 |  |  |  |
| Contrast | 23.331 | 1 | 0.000 |  |  |  |
| Accuracy | 0.786 | 1 | 0.375 |  |  |  |

The tables present the best-fitting generalized linear mixed-effects models, including fixed and random effects, degrees of freedom (df), and Akaike's Information Criterion (AIC). The model with the lowest AIC was selected as best; however, if competing models had a $\Delta AIC < 2$ , they were considered statistically indistinguishable (Burnham & Anderson, 2004), and the simpler model was preferred. In such cases, a follow-up likelihood ratio test was used to compare model fit. The final selected model is shown in italics. P < 0.05 indicates statistical significance.

**Table S7.** Summary and comparison of generalized linear mixed-effects models assessing bees’ final choice outcomes (correct, incorrect, or omission when choosing which of two rewarding chambers to enter) across varying spatial frequency trials.

| Selection of best-fit model |  |  |  |  |  |  |
| --- | --- | --- | --- | --- | --- | --- |
| Model | Dependent variable | Fixed factors | Random factor | df | AIC | ΔAIC |
| model 1 | YesNo | Treatment*Type | ID | 7 | 846.492 | 0.000 |
| model 2 | YesNo | Treatment+Type | ID | 5 | 853.253 | 6.761 |
| Type II Wald chi-square tests for fixed effects |  |  |  |  |  |  |
| Effect | X<sup>2</sup> | df | P |  |  |  |
| Treatment | 0.024 | 1 | 0.878 |  |  |  |
| Type | 328.28 | 2 | 0.000 |  |  |  |
| Treatment x Type | 9.775 | 2 | 0.008 |  |  |  |
| Bonferroni-corrected paired post hoc comparison for Treatment × Type interaction |  |  |  |  |  |  |
| Level | Estimate | SE | df | z | P |  |
| Correct | -0.518 | 0.261 | Inf | -1.986 | 0.047 |  |
| Incorrect | 0.196 | 0.29 | Inf | 0.676 | 0.499 |  |
| Omission | 1.218 | 0.524 | Inf | 2.323 | 0.02 |  |

The tables present the best-fitting generalized linear mixed-effects models, including fixed and random effects, degrees of freedom (df), and Akaike's Information Criterion (AIC). The model with the lowest AIC was selected as best; however, if competing models had a ΔAIC < 2, they were considered statistically indistinguishable (Burnham & Anderson, 2004), and the simpler model was preferred. In such cases, a follow-up likelihood ratio test was used to compare model fit. The final selected model is shown in italics. P < 0.05 indicates statistical significance

**Table S8.** Summary and comparison of generalized linear mixed-effects models assessing bees’ final choice latency during varying spatial frequency trials.

| Selection of best-fit model |  |  |  |  |  |  |
| --- | --- | --- | --- | --- | --- | --- |
| Model | Dependent variable | Fixed factors | Random factor | df | AIC | ΔAIC |
| model 1 | log(latency) | Treatment*Accuracy+Frequency | ID | 11 | 881.541 | 0 |
| model 2 | log(latency) | Treatment+Frequency+Accuracy | ID | 10 | 885.405 | 3.863 |
| model 3 | log(latency) | Treatment*Frequency*Accuracy | ID | 26 | 885.655 | 4.114 |
| model 4 | log(latency) | Treatment+Frequency*Accuracy | ID | 15 | 892.062 | 10.521 |
| model 5 | log(latency) | Treatment*Frequency+Accuracy | ID | 15 | 892.963 | 11.421 |
| Summary of model coefficients with standard errors and significance tests |  |  |  |  |  |  |
| Term | Estimate | SE | t | df | P |  |
| (Intercept) | 1.127 | 0.205 | 5.496 | 281.126 | 0.000 |  |
| Treatment Shaking | -0.009 | 0.197 | -0.045 | 143.375 | 0.964 |  |
| Correct choice | 0.171 | 0.161 | 1.063 | 315.192 | 0.289 |  |
| Frequency 0.035 deg-1 | 0.782 | 0.197 | 3.964 | 278.983 | 0.000 |  |
| Frequency 0.14 deg-1 | 1.154 | 0.17 | 6.798 | 278.082 | 0.000 |  |
| Frequency 0.28 deg-1 | 1.77 | 0.174 | 10.165 | 281.182 | 0.000 |  |
| Frequency 0.437 deg-1 | 1.915 | 0.179 | 10.719 | 283.577 | 0.000 |  |
| Frequency 0.699 deg-1 | 2.163 | 0.212 | 10.218 | 284.784 | 0.000 |  |
| Treatment Shaking:Correct choice | -0.581 | 0.217 | -2.672 | 315.028 | 0.008 |  |
| Type II Wald chi-square tests for fixed effects |  |  |  |  |  |  |
| Effect | X² | df | P |  |  |  |
| Treatment | 11.029 | 1 | 0.001 |  |  |  |
| Accuracy | 1.333 | 1 | 0.248 |  |  |  |
| Frequency | 173.049 | 5 | 0 |  |  |  |
| Treatment:Accuracy | 7.138 | 1 | 0.008 |  |  |  |
| Bonferroni-corrected paired post hoc comparison for Treatment × Type interaction |  |  |  |  |  |  |
| Level | Estimate | SE | df | z | P |  |
| Incorrect crossing | 0.009 | 0.198 | 146.359 | 0.044 | 0.965 |  |
| Correct crossing | 0.59 | 0.141 | 57.646 | 4.176 | 0 |  |

The tables present the best-fitting generalized linear mixed-effects models, including fixed and random effects, degrees of freedom (df), and Akaike's Information Criterion (AIC). The model with the lowest AIC was selected as best; however, if competing models had a $\Delta AIC < 2$ , they were considered statistically indistinguishable (Burnham & Anderson, 2004), and the simpler model was preferred. In such cases, a follow-up likelihood ratio test was used to compare model fit. The final selected model is shown in *italics*. $P < 0.05$ indicates statistical significance.

**Table S9.** Summary and comparison of generalized linear mixed-effects models assessing bee final choice accuracy on varying spatial frequency trials.

| Selection of best-fit model |  |  |  |  |  |  |
| --- | --- | --- | --- | --- | --- | --- |
| Model | Dependent variable | Fixed factors | Random factor | df | AIC | ΔAIC |
| model 1 | Accuracy | Latency*Frequency+Treatment | ID | 14 | 286.736 | 0 |
| model 2 | Accuracy | Latency+Frequency+Treatment | ID | 9 | 290.836 | 4.1 |
| model 3 | Accuracy | Latency*Treatment+Frequency | ID | 10 | 292.505 | 5.77 |
| model 4 | Accuracy | Frequency*Treatment+Latency | ID | 14 | 292.688 | 5.952 |
| model 5 | Accuracy | Latency*Frequency+Latency *Treatment+Frequency *Treatment | ID | 20 | 293.215 | 6.48 |
| Type II Wald chi-square tests for fixed effects. |  |  |  |  |  |  |
| Effect |  | X<sup>2</sup> | df | P |  |  |
| Latency |  | 5.766 | 1 | 0.016 |  |  |
| Frequency |  | 5.584 | 5 | 0.349 |  |  |
| Treatment |  | 0.009 | 1 | 0.924 |  |  |
| Latency x Frequency |  | 11.143 | 5 | 0.049 |  |  |
| Post hoc estimates of slopes for the Latency × Frequency interaction, showing the effect of latency on probability of entering incorrect reward chamber at each spatial frequency level. |  |  |  |  |  |  |
| Frequency |  | Estimate | SE | z | P |  |
| 0.07 cycles deg<sup>-1</sup> |  | -0.026 | 0.199 | -0.13 | 0.897 |  |
| 0.035 cycles deg<sup>-1</sup> |  | -0.037 | 0.017 | -2.142 | 0.032 |  |
| 0.14 cycles deg<sup>-1</sup> |  | -0.081 | 0.029 | -2.762 | 0.006 |  |
| 0.28 cycles deg<sup>-1</sup> |  | -0.01 | 0.009 | -1.007 | 0.314 |  |
| 0.437 cycles deg<sup>-1</sup> |  | -0.02 | 0.013 | -1.601 | 0.109 |  |
| 0.699 cycles deg<sup>-1</sup> |  | 0.037 | 0.025 | 1.484 | 0.138 |  |

The tables present the best-fitting generalized linear mixed-effects models, including fixed and random effects, degrees of freedom (df), and Akaike's Information Criterion (AIC). The model with the lowest AIC was selected as best; however, if competing models had a $\Delta AIC < 2$ , they were considered statistically indistinguishable (Burnham & Anderson, 2004), and the simpler model was preferred. In such cases, a follow-up likelihood ratio test was used to compare model fit. The final selected model is shown in italics. $P < 0.05$ indicates statistical significance.

